# Structure-function investigation of a new VUI-202012/01 SARS-CoV-2 variant

**DOI:** 10.1101/2021.01.01.425028

**Authors:** Jasdeep Singh, Nasreen Z. Ehtesham, Syed Asad Rahman, Seyed E. Hasnain

## Abstract

The SARS-CoV-2 (Severe Acute Respiratory Syndrome-Coronavirus) has accumulated multiple mutations during its global circulation. Recently, a new strain of SARS-CoV-2 (VUI 202012/01) had been identified leading to sudden spike in COVID-19 cases in South-East England. The strain has accumulated 23 mutations which have been linked to its immune evasion and higher transmission capabilities. Here, we have highlighted structural-function impact of crucial mutations occurring in spike (S), ORF8 and nucleocapsid (N) protein of SARS-CoV-2. Some of these mutations might confer higher fitness to SARS-CoV-2.

**Summary:** Since initial outbreak of COVID-19 in Wuhan city of central China, its causative agent; SARS-CoV-2 virus has claimed more than 1.7 million lives out of 77 million populations and still counting. As a result of global research efforts involving public-private-partnerships, more than 0.2 million complete genome sequences have been made available through Global Initiative on Sharing All Influenza Data (GISAID). Similar to previously characterized coronaviruses (CoVs), the positive-sense single-stranded RNA SARS-CoV-2 genome codes for ORF1ab non-structural proteins (nsp(s)) followed by ten or more structural/nsps [1, 2]. The structural proteins include crucial spike (S), nucleocapsid (N), membrane (M), and envelope (E) proteins. The S protein mediates initial contacts with human hosts while the E and M proteins function in viral assembly and budding. In recent reports on evolution of SARS-CoV-2, three lineage defining non-synonymous mutations; namely D614G in S protein (Clade G), G251V in ORF3a (Clade V) and L84S in ORF 8 (Clade S) were observed [2–4]. The latest pioneering works by Plante et al and Hou et al have shown that compared to ancestral strain, the ubiquitous D614G variant (clade G) of SARS-CoV-2 exhibits efficient replication in upper respiratory tract epithelial cells and transmission, thereby conferring higher fitness [5, 6]. As per latest WHO reports on COVID-19, a new strain referred as SARS-CoV-2 VUI 202012/01 (Variant Under Investigation, year 2020, month 12, variant 01) had been identified as a part of virological and epidemiological analysis, due to sudden rise in COVID-19 detected cases in South-East England [7]. Preliminary reports from UK suggested higher transmissibility (increase by 40-70%) of this strain, escalating Ro (basic reproduction number) of virus to 1.5-1.7 [7, 8]. This apparent fast spreading variant inculcates 23 mutations; 13 non-synonymous, 6 synonymous and 4 amino acid deletions [7]. In the current scenario, where immunization programs have already commenced in nations highly affected by COVID-19, advent of this new strain variant has raised concerns worldwide on its possible role in disease severity and antibody responses. The mutations also could also have significant impact on diagnostic assays owing to S gene target failures.

---

Here, we have highlighted crucial non-synonymous mutations and deletions occurring in SARS-CoV-2 VUI 202012/01 S, N and ORF8 proteins and their impact on structure-function of proteins. These proteins were selected owing to their curial role in host-viral interactome. Structural effects of SARS-CoV-2 mutations were studied using variant analysis module of COVID-3D suite [9, 10]. Images were created using PyMol [11]. In S gene, besides H69, V70 and Y144 deletion in its N-terminal domain (NTD), N501Y in the receptor binding domain (RBD), A570D in sub-domain-1 (SD1), P681H near the host processing furin cleavage site and T716I, S982A, D1118H in S2 domains were observed. The H69, V70 and Y144 were observed to be localized on solvent accessible β-hairpin loops in the NTD (Figure 1A). In case of murine coronavirus S protein, its NTD was associated with extended host range of viruses. The 69-70, 144 deletions originally observed during transmission in mink population highlights adaptation possibilities of SARS-CoV-2 in susceptible animal reservoirs [8]. In two recent studies, monoclonal antibodies isolated from convalescent COVID-19 patients were found to interact specifically with NTD of S protein and its associated mutations could confer antibody resistance [12, 13]. As per latest reports, the most concerning mutation was N501Y in RBD region can promote binding of S protein with host ACE2 receptors [14]. In crystal structure complex (PDB id: 6m0j), N501 of RBD (S protein) forms H-bond with Y41 of ACE2 receptors. Mutation analysis of the complex revealed a stronger interaction network of Y501 compared to wild-type N501 (Figure 1B). Structure based predictions showed mild stabilization of the mutant (ΔΔG^stability^ SDM~ 0.4 kcal.mol^-1^, Table 1). The N501Y mutation was, however distantly located from CR3022 and C135 (SARS-CoV-2 neutralizing monoclonal antibodies) and RBD interaction interface (Supplementary Figure 1 and Supplementary Figure 2A) [15]. However, both of these two antibodies share distinct epitopes on RBD of S protein (Supplementary Figure 2B). A recent analysis (preprint) identified new genetic variants of SARS-CoV-2 which can evade host immune responses [16]. In Australia, out of 24 immune escape variants, S477N mutant was present in >60% of genome samples. This mutation was located outside RBD-C135/CR3022 interface (Supplementary Figure 2B). For India, N440K high frequency variant was observed out of 19 immune escape variants [16]. This mutation is located at C135 interaction interface. In wild type strain, N440 forms a strong H-bond network with D54 and weak H-bond networks with P52 and R55 of C135 antibody (Supplementary Figure 2C). The K440 mutant however, only forms a weak H-bond with D54 (Supplementary Figure 2D). This could possibly explain immune evasion as suggested by Gupta et al for asymptomatic reinfection of this mutant in two healthcare workers from India [17]. Interestingly, we observed 100% co-occurrence on N440K mutation along C64F mutation in membrane glycoprotein of SARS-CoV-2 apart from globally dominant D614G (S protein) and P323L (ORF1ab) [4, 9]. The other new mutations in S protein showed co-occurrence with only D614G-P323L type [9]. Taken together, the naturally occurring mutations might confer resistance against one antibody, not other.

**Figure 1.**
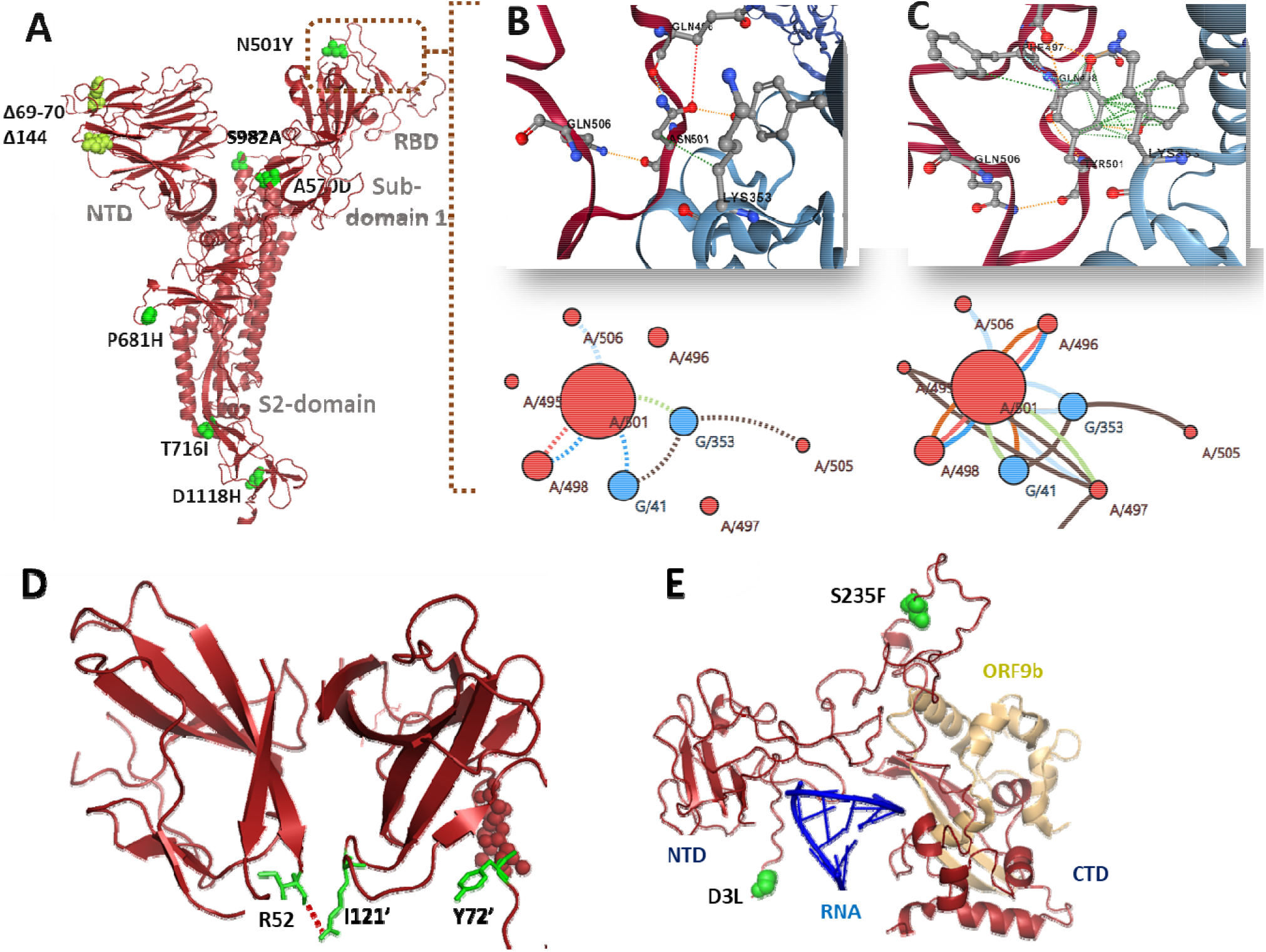
Structural localization of mutations in S, ORF8 and N proteins of SARS-CoV-2 VUI-202012/01 variant. (A) Cartoon representation of S protein protomer, showing non-synonymous mutations (green spheres) and deletions (yellow-green spheres) in VUI-202012/01 variant. The N501Y mutation occurs in the RBD, which forms first contact with host ACE receptors. (B) Residue interaction network at RBD (Brown)-ACE2 (Blue) interface for wild type N501 variant. Panel below shows detailed interaction of N501 with other residues of RBD of S protein (marked as Chain A) and ACE2 (marked as Chain G) (Dashed lines). (C) Residue interaction network at RBD (Brown)-ACE2 (Blue) interface for mutant Y501 variant. Panel below shows detailed interaction of Y501 with other residues of RBD of S protein (marked as Chain A) and ACE2 (marked as Chain G) (Straight lines). Color codes: H-bonds (red), Polar H-bonds (orange), VdW (light blue), Aromatic (light green) and Ring-ring interactions (brown). (D) Cartoon representation of ORF8 (brown) dimer showing H-bond interactions of R52 and I121’ (green sticks) between two monomers and Y72’. (E) Cartoon representation of N protein RNA (blue) interacting NTD and ORF9b (golden yellow) interacting CTD. The two D3L and S235F mutations are localized in the unstructured NTD and linker regions, respectively.

The ORF8 protein of SARS-CoV-2 is proposed to interact with variety of host proteins and modulate immune responses; and disruption of IFN-I signaling and downregulation of MHC-I in cells [18, 19]. Crystal structure of ORF8 showed two dimerization interfaces unique to SARS-CoV-2 genealogy. The R52I and Y73C mutations in SARS-CoV-2 VUI 202012/01 are localized at these interaction interfaces. In wild type strain, R52 forms H-bond contact with I121 from another protomer (Figure 1D). Y73 forms part of crucial _73_YIDI_76_ non-covalent ORF8 dimer interaction interface, unique to SARS-CoV-2. Presence of these interfaces enables ORF8 to form large scale assemblies, possibly aiding SARS-CoV-2 to evade and modulate host immune responses [18].

The SARS-CoV-2 N protein can exist both in monomeric and oligomeric forms, and interact with RNA. The dimerization interface is formed through interactions of C-terminal domain (256-364, PDB id: 6wzo) while the RNA interactions are mediated through its N-terminal domain (46-176, PDB id: 7acs) (Figure 1E). Recently, it was shown that ORF9b (alternative ORF within N gene) can suppress IFN-I responses through physical interactions with mitochondrial TOM70 or induction of lactic acid production [20]. The D3L and S235F mutations in the N protein occur outside the dimer and RNA interaction interfaces, in the unstructured regions in its NTD and linker regions, respectively (Figure 1E). However, mutation analysis predicted stabilization effect conferred by S235F to N protein (Table 1).

**Table 1:**
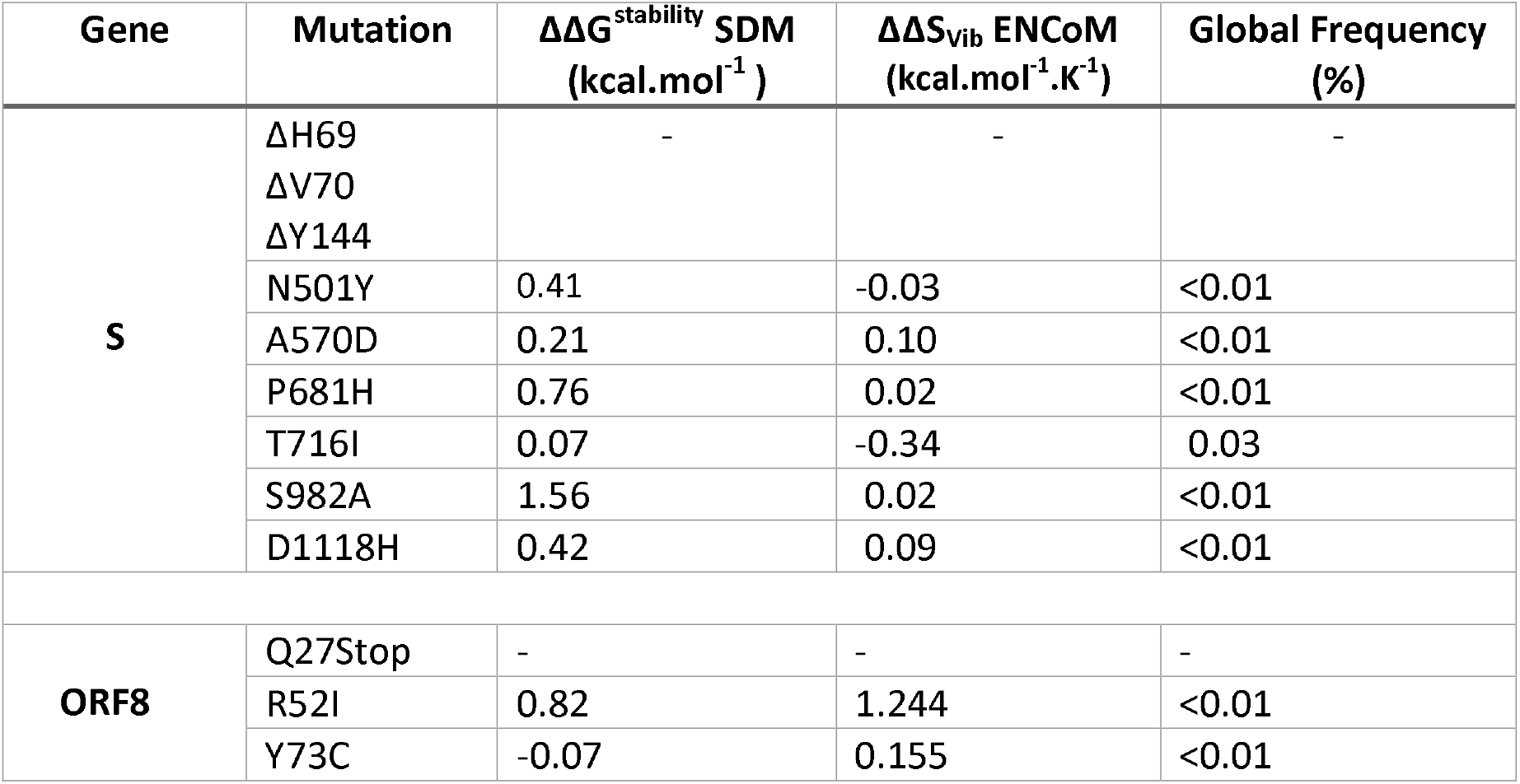
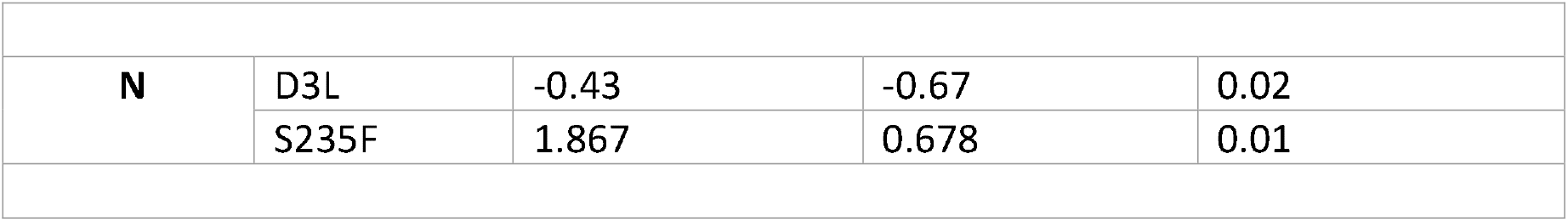
Notable non-synonymous mutations and deletions in S, ORF8 and N proteins of SARS-CoV-2 VUI-202012/01 variant. DynaMut analysis for predicting impact of mutations of structural stability of proteins. Positive ΔΔG^stability^ SDM and ΔΔS_Vib_ ENCoM values indicate stabilizing mutations with increase in molecule flexibility. Negative values indicate destabilizing mutations with decrease in molecule flexibility. Table adapted from Portelli et al and Public Health England report [9, 14].

| Gene | Mutation | $\Delta\Delta G^{\text{stability}}$ SDM<br>(kcal.mol<sup>-1</sup>) | $\Delta\Delta S_{\text{vib}}$ ENCoM<br>(kcal.mol<sup>-1</sup>.K<sup>-1</sup>) | Global Frequency<br>(%) |
| --- | --- | --- | --- | --- |
| **S** | ΔH69<br>ΔV70<br>ΔY144 | - | - | - |
|  | N501Y | 0.41 | -0.03 | <0.01 |
|  | A570D | 0.21 | 0.10 | <0.01 |
|  | P681H | 0.76 | 0.02 | <0.01 |
|  | T716I | 0.07 | -0.34 | 0.03 |
|  | S982A | 1.56 | 0.02 | <0.01 |
|  | D1118H | 0.42 | 0.09 | <0.01 |
| **ORF8** | Q27Stop | - | - | - |
|  | R52I | 0.82 | 1.244 | <0.01 |
|  | Y73C | -0.07 | 0.155 | <0.01 |
| N | D3L | -0.43 | -0.67 | 0.02 |
|  | S235F | 1.867 | 0.678 | 0.01 |

In conclusions, mutations in S and ORF8 proteins can have significant impact on viral transmission and host immune responses. Coupling of viral pathogenesis and clinical data with long range molecular dynamics simulations can further aid in understanding structure function impacts of these mutations.

---

**Supplementary Figure 1.**
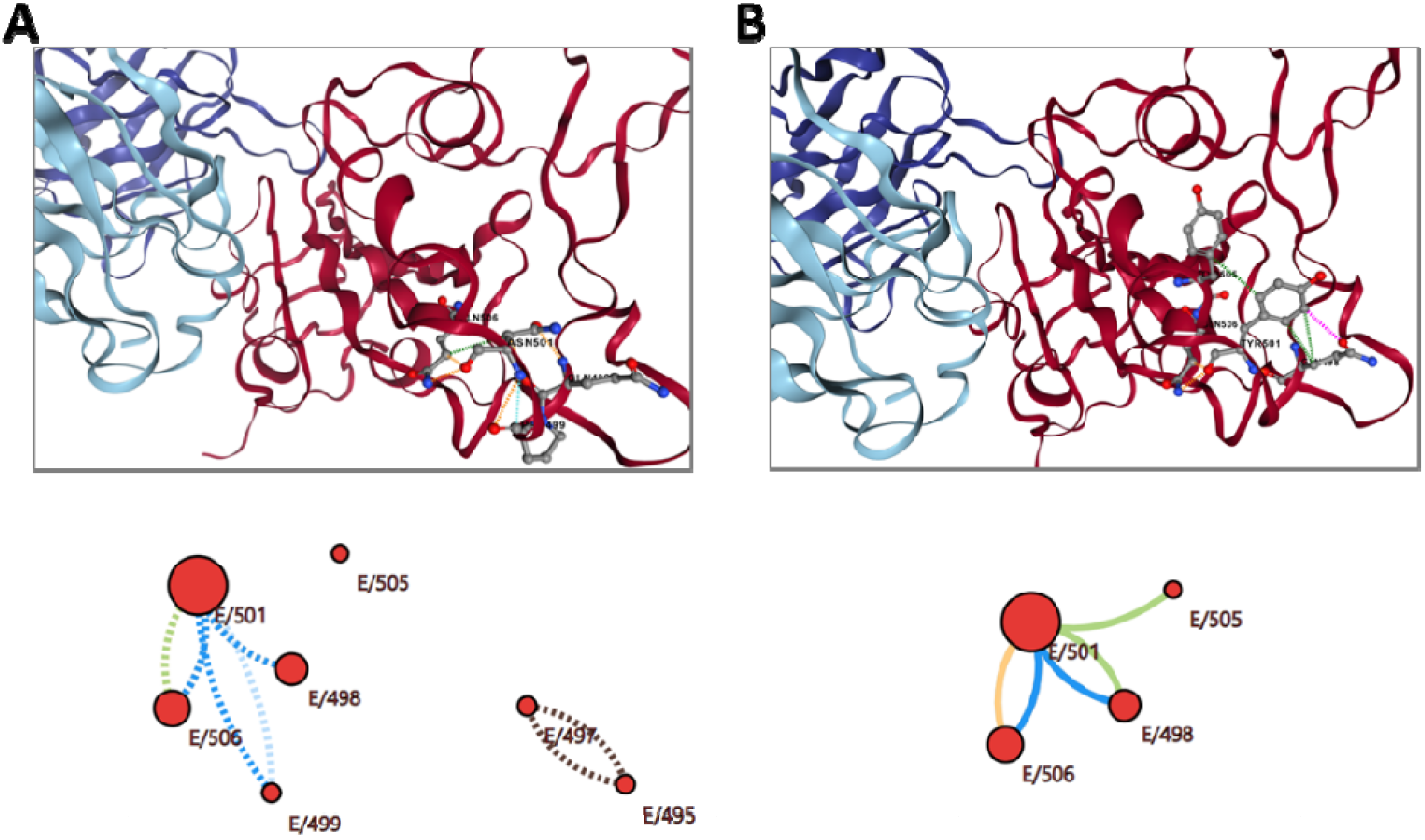
(A) Residue interaction network at RBD (Brown)-CR3022 (SARS-CoV-2 neutralizing monoclonal antibody, Blue) interface for wild type N501 variant. Panel below shows detailed interaction of N501 with other residues of RBD of S protein (marked as Chain E, Dashed lines). (B) Residue interaction network at RBD (Brown)- CR3022 (Blue) interface for mutant Y501 variant. Panel below shows detailed interaction of Y501 with other residues of RBD of S protein (marked as Chain A, straight lines). Color codes: H-bonds (red), Polar H-bonds (orange), VdW (light blue), Aromatic (light green) and Ring-ring interactions (brown). N501 or Y501 is not involved in direct interaction with CR3022 antibody.

**Supplementary Figure 2.**
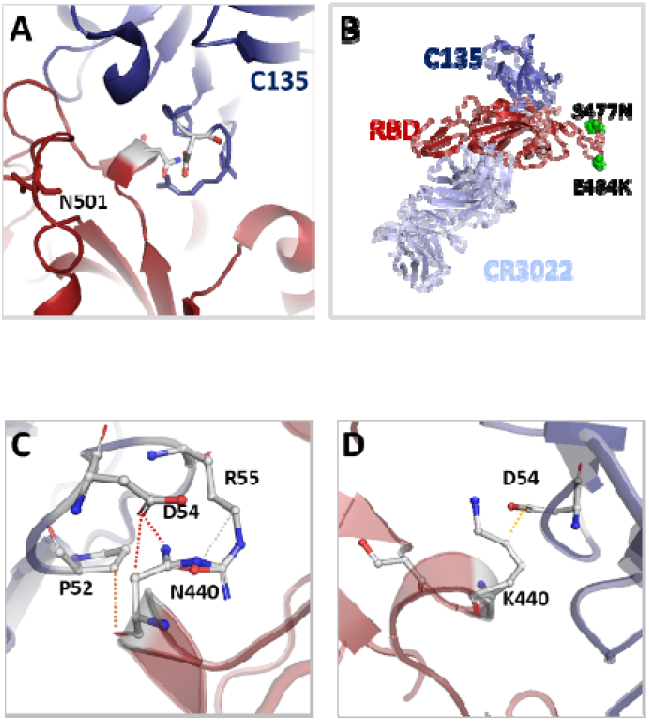
(A) Interaction interface between RBD (Brown) of S protein of SARS-CoV-2 and human monoclonal C135 anti-SARS-CoV-2 antibody (Blue). (B) Distinct interaction interface (epitope) between RBD (Brown) and C135 (Dark Blue) and CR0322 (Light Blue) antibodies. S477N and E484K mutations occur outside both RBD-antibody interfaces (shown as green spheres). (C) Residue interaction network of N440 (RBD, Brown) with residues of C135 antibody (Blue). (D) Residue interaction network of K440 (RBD, Brown) with residues of C135 antibody (Blue). Strong H-bonds are shown as red and weak H-bonds as orange.

